# Long-term surveillance defines spatial and temporal patterns implicating *Culex tarsalis* as the primary vector of West Nile virus in Iowa, USA

**DOI:** 10.1101/476234

**Authors:** Brendan M. Dunphy, Kristofer B. Kovach, Ella J. Gehrke, Eleanor N. Field, Wayne A. Rowley, Lyric C. Bartholomay, Ryan C. Smith

## Abstract

West Nile virus (WNV) has become the most epidemiologically important mosquito-borne disease in the United States, causing ∼50,000 cases since its introduction in 1999. Transmitted primarily by *Culex* species, WNV transmission requires the complex interplay between bird reservoirs and mosquito vectors, with human cases the result of epizootic spillover. To better understand the intrinsic factors that drive these interactions, we have compiled infection data from sentinel chickens, mosquito vectors, and human cases in Iowa over a 15 year period (2002-2016) to better understand the spatial and temporal components that drive WNV transmission. Supplementing these findings with mosquito abundance, distribution, and host preferences data, we provide strong support that *Culex tarsalis* is the most important vector of human WNV infections in the region. Finally, we identify underlying climatic factors (temperature and drought) that are associated with inter-annual trends in WNV intensity. Together, our analysis provides new insights into WNV infection patterns in multiple hosts and highlights the importance of long-term surveillance to understand the dynamics of mosquito-borne-disease transmission.

## Introduction

In an era of increased concern over mosquito-borne viruses, West Nile virus (WNV) continues to have the largest epidemiological impact in the United States, causing ∼50,000 cases and over 2,100 deaths since its introduction in 1999 (1). Therefore, the annual occurrence of WNV presents a continual public health threat across the continent.

WNV persists in nature through a host cycle that involves bird reservoirs and *Culex* mosquito vectors, with humans serving as dead-end hosts through the bite of an infected mosquito (2). Both abiotic factors (e.g.-landscape, climate, and seasonality) and biotic factors (e.g.-community composition of bird reservoirs and mosquito vectors, species abundance, and inter-species interactions) influence WNV infection dynamics across the United States (3–7). Human cases of WNV have been reported across all 48 states of the contiguous US, yet specific states or geographic regions have been impacted by a disproportionate number of cases (1). Three states (California, Colorado, and Texas) comprise more than 36% of the total number of reported WNV cases, while states of the Upper Midwest, particularly South Dakota and Nebraska, have consistently displayed among the highest incidence rates (cases per capita) in the country (1). As a result, it is of great importance to understand the epidemiological patterns of WNV across the country (8).

Previous reports describing patterns of WNV transmission in densely populated urban/suburban areas (9–12) and rural environments (13, 14), suggest that distinct mechanisms of peridomestic and sylvatic transmission may influence the epidemiology of WNV across the US (8). This is supported by considerations of land use and landscape ecology, which serve as important determinants in shaping the geographical distributions of mosquito vectors (15–19). Of the *Culex* species implicated as vectors of WNV, *Culex pipiens, Culex quinquefasciatus*, and *Culex restuans* are routinely associated with urbanized settings (20, 21) while *Culex tarsalis* is most common in rural habitats (15, 16, 22). With known differences in vector competence (23, 24), the spatial distributions of these *Culex* species can have profound impacts on patterns of WNV transmission at the local or regional level. Factors influencing WNV transmission have been broadly described across the United States (8, 25–27), yet our understanding of WNV transmission in the Midwest has heavily relied on studies of the Chicago metropolitan area (12, 28–33) with only limited characterization of WNV epidemiology and transmission dynamics in other locations of the Midwest (14, 15, 34–36).

Building on initial reports describing WNV transmission in Iowa from 2002-2006 (15), we have assembled an extensive 15 year study of WNV transmission for the state (2002 - 2016) examining WNV infection data from human, sentinel chicken, and mosquito hosts. Combined with mosquito abundance, distribution, and host selection data, our comprehensive resources offer the unique ability to investigate the concurrent impacts of multiple ecological factors on human WNV disease cases. By examining host infection rates from spatial and temporal perspectives, we have determined when and where WNV is most actively being transmitted in Iowa, which suggests underlying biotic and abiotic mechanisms that likely influence WNV transmission. Together, these data provide strong support that *Cx. tarsalis* is the predominant vector of human WNV transmission in Iowa. These analyses provide new insights into WNV transmission dynamics in the Upper Midwest region and highlight the importance of long-term surveillance to understand mosquito borne-disease.

## Materials & Methods

### West Nile virus surveillance program

Since 2002, the Medical Entomology Lab at Iowa State University has conducted WNV surveillance for the state of Iowa through cooperative efforts with the Iowa Department of Public Health, the State Hygienic Lab, and city- or county-level municipalities around the state, where field operations occurred. Data presented herein were collected over the 15 year period from 2002 to 2016, generally between mid-May and early October (approximately weeks 20 - 42) of each year as the period of mosquito activity in Iowa.

### Mosquito collections and identification

Adult mosquitoes were collected across the state using a variety of trapping methods to assess population dynamics or to monitor WNV infection rates. New Jersey light traps (NJLTs) were used to measure mosquito abundance, while CO2-baited CDC light traps (CDC traps) or grass infusion-baited gravid traps were used to collect mosquitoes for subsequent WNV testing. All three trap types were used for the duration of the 15 year study period, with the exception of a two year period in which CDC trapping did not occur (2014 and 2015) and one year (2002) in which gravid traps were not used. Additional trapping methods relying on Mosquito Magnets (2007–2009) or BG-Sentinel traps (2016) were selectively used across the state. All traps were maintained and operated by either Iowa State University personnel or local municipal collaborators.

Mosquito samples were identified according to morphological characteristics described for North American mosquitoes (37). To keep data consistent across the 15 year trapping period and the difficulty in distinguishing adult female *Cx. pipiens, Cx. quinquefasciatus, Cx. restuans*, and *Cx. salinarius* (especially when samples are in poor condition), these species were collectively identified as “*Cx. pipiens* group” due to the difficulty in distinguishing these species as adults, especially in the large volumes processed in our study with specimens frequently in poor physical condition (38, 39). This nomenclature was used throughout the study.

### West Nile virus (WNV) testing in mosquitoes

Mosquito samples collected from CDC and gravid traps were routinely used for WNV testing during the 15 year sample period. Additional samples collected from BG-Sentinel and Mosquito Magnet traps were respectively tested for only a single season or a small period of 3 years. Following collection, mosquitoes were stored at either -20 °C or −80°C by our collaborators before shipping in insulated parcels with frozen ice packs or dry ice. To preserve samples during identification and handling, mosquitoes were sorted on refrigerated tabletops. After identification, mosquitoes of the same species were grouped together into pools not exceeding 50 mosquitoes according to trap type, trap site and collection week. Mosquito pools were sent to the State Hygienic Laboratory (Iowa City, IA) for WNV detection by quantitative RT-PCR. Only data collected for *Culex* species that regularly tested positive for WNV (i.e., *Cx. pipiens* group, *Cx. tarsalis*, and *Cx. erraticus)* were used in this study and were collectively referred to as “*Culex*” species (Table S1).

Mosquito infection rates were calculated as minimum infection rates (MIR) from bias-corrected maximum likelihood estimates using the CDC-provided Microsoft Excel add-in for calculation of pooled infection rates (40). Mosquito infection data were parsed according to week of the year, mosquito species, trap type, and county of collection.

### Sentinel chicken seroprevalence

Sentinel chickens were used in participating Iowa municipalities from 2002 to 2013 to monitor WNV seroprevalence in local wild bird reservoirs. At the beginning of each surveillance year, baseline bleeds were performed on each chicken to ensure the absence of WNV infection. Sentinel “flocks” consisting of 8 chickens were deployed to sites around the state, based on the availability of municipal collaborators to maintain the flocks. Weekly blood draws from each individual chicken (target volume – 0.5 mL) were taken to gauge infection status through the summer. All blood samples were sent to the State Hygienic Laboratory (Iowa City, IA) to test for the detection of IgM antibodies to WNV using a MAC-ELISA assay as previously described (41). Seroprevalence was calculated by dividing the number of infected chickens by the total number of sampled chickens in a population. Animal procedures were approved by the Iowa State University Institutional Animal Care and Use Committee protocol #7-2-5196G to L. Bartholomay.

### Human case reporting

As a mandatory reportable disease, human cases of WNV were reported to the Iowa Department of Public Health from physicians and health clinics around the state, before forwarding information to the Centers for Disease Control (CDC). All human case data were reported at the county level by week of the year (except 2002, which lacked temporal data). Human incidence was calculated by dividing the number of reported human cases by the human population (Iowa county or region) and normalized per 100,000 people.

### Mosquito blood meal analysis

Mosquitoes containing a visible blood meal that were collected in CDC light traps or gravid traps at various locations throughout the state were preserved at −80°C. This included all *Cx. tarsalis* blood-fed samples in our possession collected from 2007 to 2011 and in 2017, a total of 93 samples. A selection of 214 blood-fed *Cx. pipiens* group samples collected from 2015 and 2017 were similarly processed. DNA was isolated from individual mosquitoes using the Marriot DNA extraction procedure (42). DNA from the Cx. tarsalis samples were validated using actin primers (43), while *Cx. pipiens* group samples were speciated by PCR analyses as previously (44) to confirm their identification as *Culex restuans, Culex pipiens*, or *Culex salinarius*. Positive DNA amplification was confirmed in 92 of 93 *Cx. tarsalis* and 159 of 214 *Cx. pipiens* group samples.

To determine the vertebrate host source of the mosquito blood meals, two methods were employed. For *Cx. tarsalis*, a PCR-RFLP assay in which cytochrome B (cytB) sequences were amplified by PCR, then digested with *Taq*I to distinguish between vertebrate species was performed on the samples (45). However, due to similarities in the *Culex pipiens* group cytB sequences with the associated PCR-RFLP primers, this assay could not distinguish vertebrate blood samples. As a result, the *Culex pipiens* group samples were evaluated by PCR using individual avian, mammal, and human primer sets as previously (31, 46). PCR reactions were performed using DreamTaq (Thermo Fisher) and visualized by electrophoresis on 1.5% agarose gels.

### Mapping of WNV infection data and mosquito abundance data

To visualize host (human, chicken, and mosquito) infection rates and mosquito abundance on a geographic scale, county-specific values were mapped using ArcMap 10.4.1 (Esri ArcGIS). Values of human incidence, sentinel chicken seroprevalence, mosquito MIR, and mosquito abundance ratios were interpolated between counties according to the inverse distance weighted (IDW) method using centroid locations of the corresponding counties as previously described (47). To enhance the visual display, a 2% clip and gamma stretch were applied.

### Statistical analyses

All variables examined in this study (host infection rates by week, year, region, and county; county-level mosquito abundance measures) were found to have non-normal distributions, with the exception of regional mosquito MIR differences. As a result, non-parametric methods were chosen for all statistical analyses to maintain consistency. Significance of relationships was determined with a 95% confidence interval (*P* < 0.05, two-tailed). A Wilcoxon signed-rank test was performed to determine differences in annual host infection rates between Iowa regions and between *Culex* species. Data were analyzed and prepared using GraphPad Prism 6 (GraphPad Software).

## Results

### WNV displays defined temporal patterns in its human, avian, and mosquito hosts

To better understand the seasonality of WNV transmission in Iowa, fifteen years of WNV surveillance data were analyzed from the three major host species (humans, birds, and mosquitoes) to identify peak periods of WNV activity. Between mid-May and early October (approximately weeks 20-42), mean weekly WNV activity is displayed for human cases (Fig. 1A), chicken seroprevalence (Fig.1B), and mosquito infection rates (Fig. 1C). Across host species, WNV activity followed similar temporal patterns reaching detectable levels approximately in early June (week 23) and amplifying progressively through the summer months. Human and mosquito infection rates peaked in early September (humans, week 36; mosquito, week 37), then rapidly declined to barely detectable levels in October (week 40). In contrast, chicken seroprevalence continued to rise through late September and early October when human incidence and mosquito infection rates have already rapidly declined (Fig. 1). Seroprevalence was not measured beyond this point as it marked the end of the field season.

**Figure 1.**
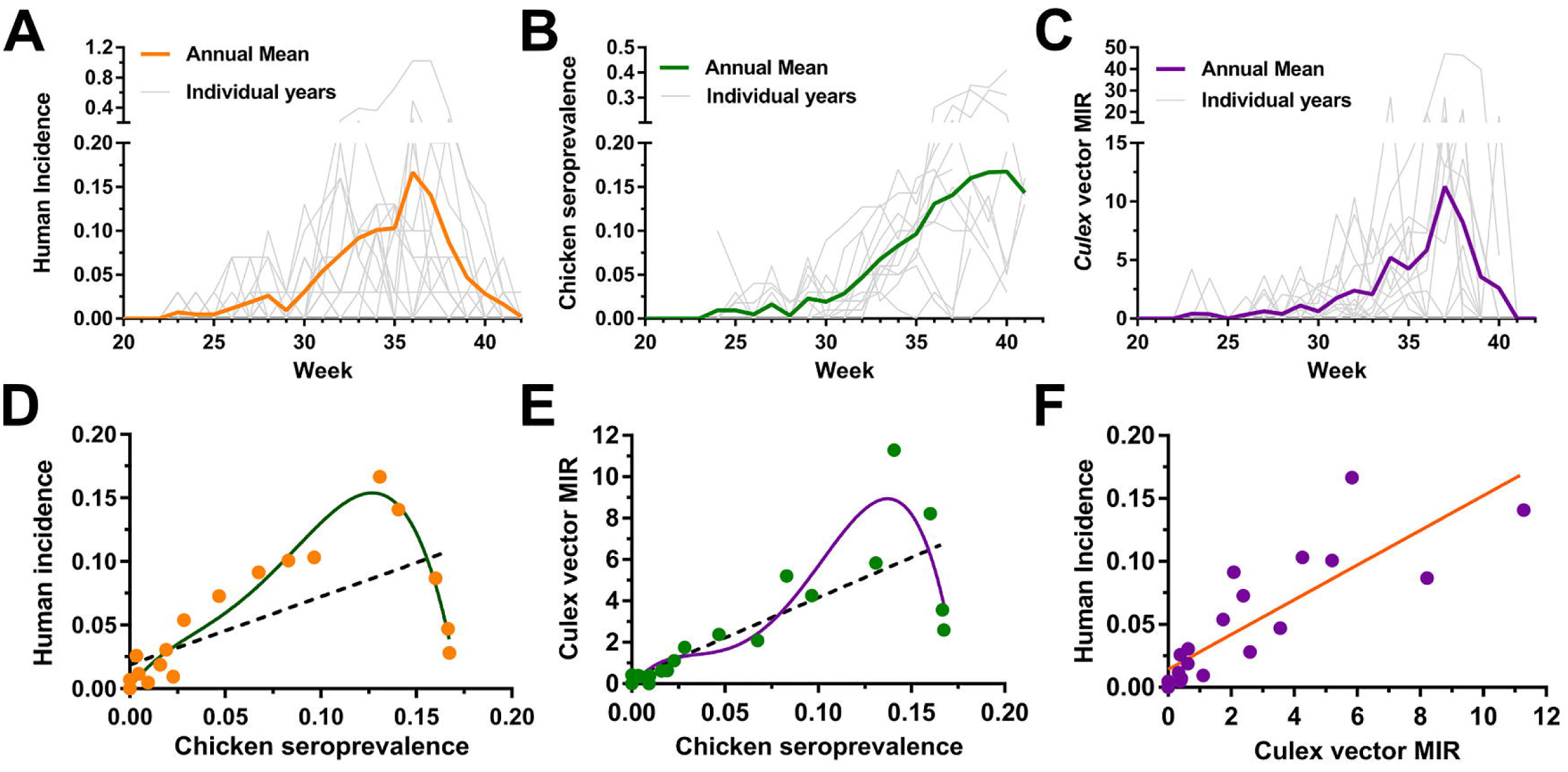
Seasonality of WNV transmission in human, avian, and mosquito hosts. WNV infection data from Iowa (2002-2016) were analyzed and displayed as weekly values for individual years or as the annual mean of human incidence (A), sentinel chicken seroprevalence (B), or *Culex* minimum infection rate (MIR) (C). Correlations between different host (human, avian, mosquito) infection rates were examined by plotting averaged weekly values (weeks 20-40; data points on graph) to determine the strength of host interactions involved in WNV transmission (D-F). A linear regression (black dashed line; R^2^=0.45, *P* <0.001) or a best-fit 4^th^ order polynomial (green line; R^2^=0.94, *P* <0.0001) were used to explain sentinel chicken seroprevalence correlations to human incidence (D). Similar linear regression (black dashed line; R^2^=0.63, *P*<0.001) and best-fit 4^th^ order polynomial (purple line; R^2^=0.86, *P* <0.0001) were performed to examine the effects of *Culex* MIR in estimating sentinel chicken seroprevalence (E). The non-linear relationship best explains the declining human incidence or mosquito MIR, while sentinel chicken seroprevalence remained high at the end of the season (E). Linear regression (orange line; R^2^=0.71, *P* <0.0001) analysis of *Culex* MIR and human WNV incidence (F).

Relationships between host infection patterns were more closely examined using regression analyses of historical infection rates (weeks 20-40) to determine the strength of these multivariate interactions. A linear model of chicken seroprevalence was limited in its ability to account for variation in human incidence, yet was greatly improved when examined as a non-linear relationship to account for the late-season decline in human incidence while seroprevalence continued to increase (Fig. 1D). Similarly, a non-linear analysis improved the ability of sentinel chicken seroprevalence to estimate variation in *Culex* minimum infection rates (MIR) (Fig. 1E). Lastly, *Culex* MIR accounted for 71% of the seasonal variation in human incidence when examined as a linear relationship (Fig.1F). Together, these data suggest that both linear and non-linear relationships of host infection rate data account for the dynamic nature of WNV activity. Traditionally used as tools for arbovirus surveillance (48, 49), these data provide strong support for the continued use of sentinel chickens and the use of mosquito infection rates to actively monitor WNV activity.

### Regional differences in WNV transmission

WNV infections were then examined across the state to identify spatial patterns of endemicity and regions of higher risks for human infection. To approach this question, mean annual infections rates were mapped for each host species (human, chicken, mosquito) at the county level for all locations in which data have been collected from 2002 to 2016 (Fig. 2). Annual infection rates were also broken down by region, with additional comparisons of infection intensity with respect to time (Fig. 2). From these data, the human incidence of WNV was determined to be the highest in western Iowa, where infection rates were four times that of central Iowa and seven times that of eastern Iowa (Fig. 2A). Central Iowa also displayed significantly higher human incidence than in eastern Iowa (Fig. 2A). Temporal analyses of human WNV incidence further support this observation, demonstrating that incidence rates in western Iowa were higher and amplified faster than in other geographic regions of the state (Fig. 2A). These data confirm previous reports of WNV incidence in Iowa from 2002 to 2006 (15), which similarly define a distinct gradient where the highest incidence rates are found in western Iowa and the lowest incidence in eastern Iowa. This regional dichotomy suggests that important entomological and ecological differences across the state must influence higher rates of enzootic spillover of WNV in western Iowa.

**Figure 2.**
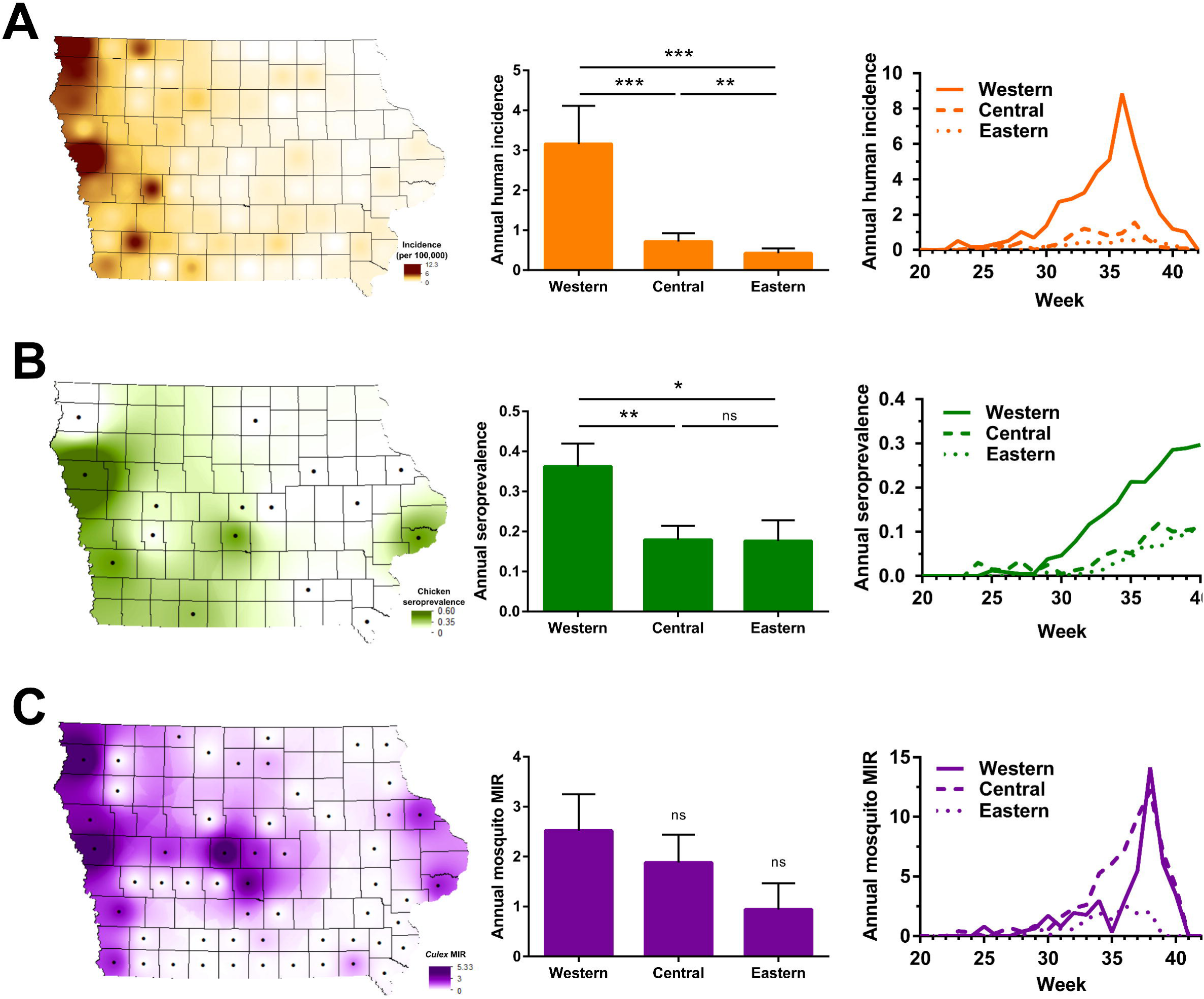
Spatial distributions of WNV in human, avian, and mosquito hosts. WNV infections from 2002-2016 are displayed at the county level and spatially presented as the annual mean of human incidence (A), sentinel chicken seroprevalence (B), or *Culex* MIR (C) across the state. Additional analyses display the annual mean or weekly infection rates by regional associations in Iowa – western, central, and eastern (A-C). Human incidence (A) is displayed as the number of human WNV cases per 100,000 people. Sentinel chicken seroprevalence (B) is defined as the ratio of WNV+ seropositive birds per total birds sampled. *Culex* MIR (C) is calculated as the minimum infection rate for all *Culex* mosquito vectors (*Cx. pipiens* group, *Cx. tarsalis*, and *Cx. erraticus*). The black dots display counties in which chicken seroprevalence and mosquito infection rate data were collected. A Wilcoxon matched-pairs signed rank test was used to test for differences between groups with significance denoted by asterisks (*P<0.05, **P<0.01, ***P<0.001). ns, not significant.

Seroconversion rates in sentinel chickens were also highest in sentinel chickens in western Iowa, displaying twice the seroconversation rates than those in central and eastern Iowa (Fig.2B). However, unlike human incidence, these hotspots were less defined with several non-western counties exhibiting high infection rates (Fig. 2B). Seroconversion in western Iowa occurred earlier in the summer, resulting in infection rates that were three times greater than those found in other regions by the end of the year (Fig. 2B). No differences in seroconversion were detected between central and eastern Iowa (Fig. 2B).

With similar spatial patterns of WNV in humans and sentinel chickens, we expected that the MIR of *Culex* species producing WNV+ mosquito pools (Table S1) would likewise display a similar pattern across the state. Although the majority of counties with the highest infection rates were found in western Iowa, we found that the overall geographic trends in mosquito infection rates were less pronounced (Fig. 2C). Mosquitoes from western Iowa displayed a greater infection rate than those from central or eastern regions, yet these differences were not significant (Fig. 2C). Temporal analyses displayed equivalent amplification of mosquito infections in western and central Iowa, with much lower mosquito infection rates in the east throughout the year (Fig. 2C). While consistent with the geographic gradients for human incidence and chicken seroprevalence, these mosquito infection trends argue that a more detailed analysis of mosquito species composition on infection rates may be required to delineate potential differences between *Culex* species as vectors of WNV transmission.

### *Cx. tarsalis* display higher infection rates than *Cx. pipiens* group mosquitoes

Since the introduction of WNV into Iowa in 2002, *Cx. pipiens* group mosquitoes and *Cx. tarsalis* have comprised 98% of the WNV+ mosquito pools (Table S1). To better understand the contributions of these vectors to WNV transmission, we examined mosquito infection data obtained from mosquitoes collected with gravid or CO2-baited CDC traps. Gravid traps lure mosquitoes seeking a site for oviposition and have been widely used for monitoring populations of *Cx. pipiens* group species (50). Likewise, mosquito yields from gravid traps were severely biased towards *Cx. pipiens* group (99.5% of 93,456 collected; Fig. 3A). However, despite the sparsity of *Cx. tarsalis* data from gravid trap collections, *Cx. tarsalis* displayed higher infection rates than *Cx. pipiens* group mosquitoes (Fig. 3B).

**Figure 3.**
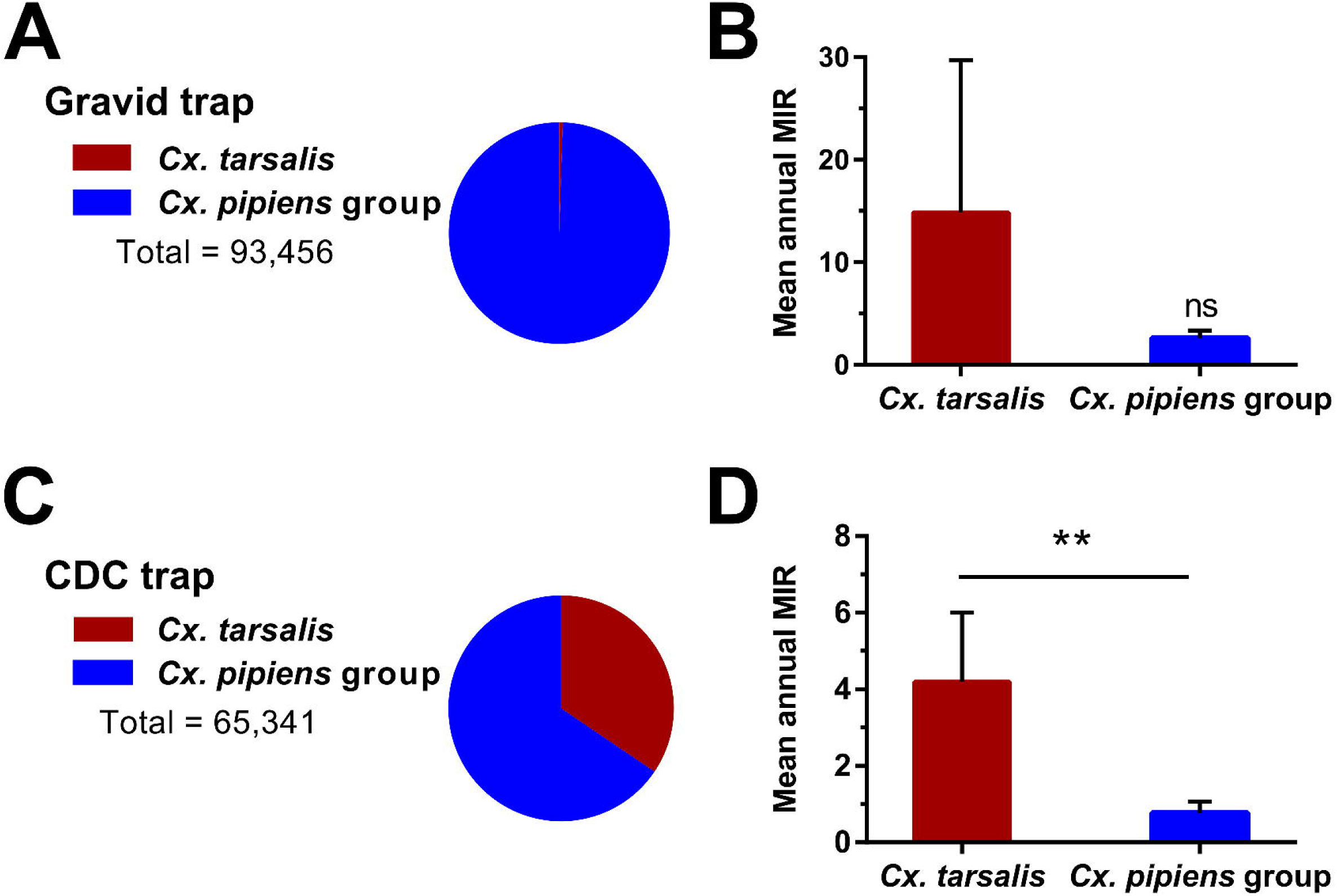
WNV surveillance reveals mosquito trapping bias and greater infection rates in *Cx. tarsalis*. (A) Gravid trap yields of *Cx. tarsalis* (red) and *Cx. pipiens* group (blue) show significant trap bias toward collecting container-breeding *Cx. pipiens* group mosquitoes. Mean annual minimum infection rates (MIR) of *Cx. tarsalis* and *Cx. pipiens* group collected from gravid traps (B). Only two years of substantial *Cx. tarsalis* yields were collected as a result of this trapping bias, resulting in a large margin of error (B). In contrast, CDC trap yields of *Cx. tarsalis* (red) and *Cx. pipiens* group (blue) showed a more even sampling of the two vectors (C), in which the mean annual MIR of *Cx. tarsalis* were significantly higher in CDC trap collections (D). The Wilcoxon matched-pairs signed test was used to test for differences between groups. Significant differences are indicated by asterisks (**P<0.01). ns, not significant.

Generally targeting mosquitoes seeking a blood meal (50), CO2-baited CDC traps resulted in more comparable yields of *Cx. tarsalis* and *Cx. pipiens* group mosquitoes (Fig. 3C). *Cx. pipiens* group were still the more abundant vector (65.5% of 65,341 collected; Fig. 3C), but these data do not account for potential differences in species distributions throughout the state. Comparisons of mean annual MIR between mosquito species revealed that the infection rate in *Cx. tarsalis* was significantly higher (∼5x) than that of *Cx. pipiens* group mosquitoes (Fig. 3D). Therefore, *Cx. tarsalis* populations in Iowa displayed higher infection rates than the *Cx. pipiens* group population during the entire 15 year study period. This is further supported by laboratory and field-based studies arguing that *Cx. tarsalis* is one of the most competent vectors of WNV in North America (24, 51, 52).

### Blood meal analysis reveals differences in host selection between *Culex* vectors

To examine potential differences in host selection between *Cx. tarsalis* and *Cx. pipiens* group, we used PCR-based methods to analyze mosquito blood meals (30, 46, 53). Of the mosquito samples analyzed, only a small percentage resulted in blood meal identifications using our PCR-based methods (25% of *Cx. tarsalis* [23 of 92] and 20% of *Cx. pipiens* group [32 of 159] samples). This is likely due to the passive collection of these blood-fed mosquito samples in which host DNA in the blood meals may have been digested past the point of PCR identification (54). These limitations prevent us from making any strong assertions about differences in host preference between these mosquito vectors. However, our data show that *Cx. tarsalis* feeds predominantly on birds (∼52% of all samples) and humans accounted for a large component (∼30%) of the identified blood meals (Fig. 4). This is in contrast to *Cx. pipiens* group mosquitoes that collectively displayed stronger selection for non-human mammals and birds, with humans representing the smallest proportion (∼13%) of analyzed samples (Fig. 4). When further classified by species using a multiplex PCR assay (44), we saw notable differences within the *Cx. pipiens* group. *Cx. restuans* (which comprised the majority of samples in our analysis) and *Cx. salinarius* fed mostly on non-human mammals (Fig. S1), while *Cx. pipiens* samples fed primarily on birds (Fig. S1). Together, these data suggest that *Cx. tarsalis* may exhibit different preferences for host selection than *Cx. pipiens* group and feed more prevalently on humans (Fig. 4), providing support that *Cx. tarsalis* may serve as a better vector of WNV to humans.

**Figure 4.**
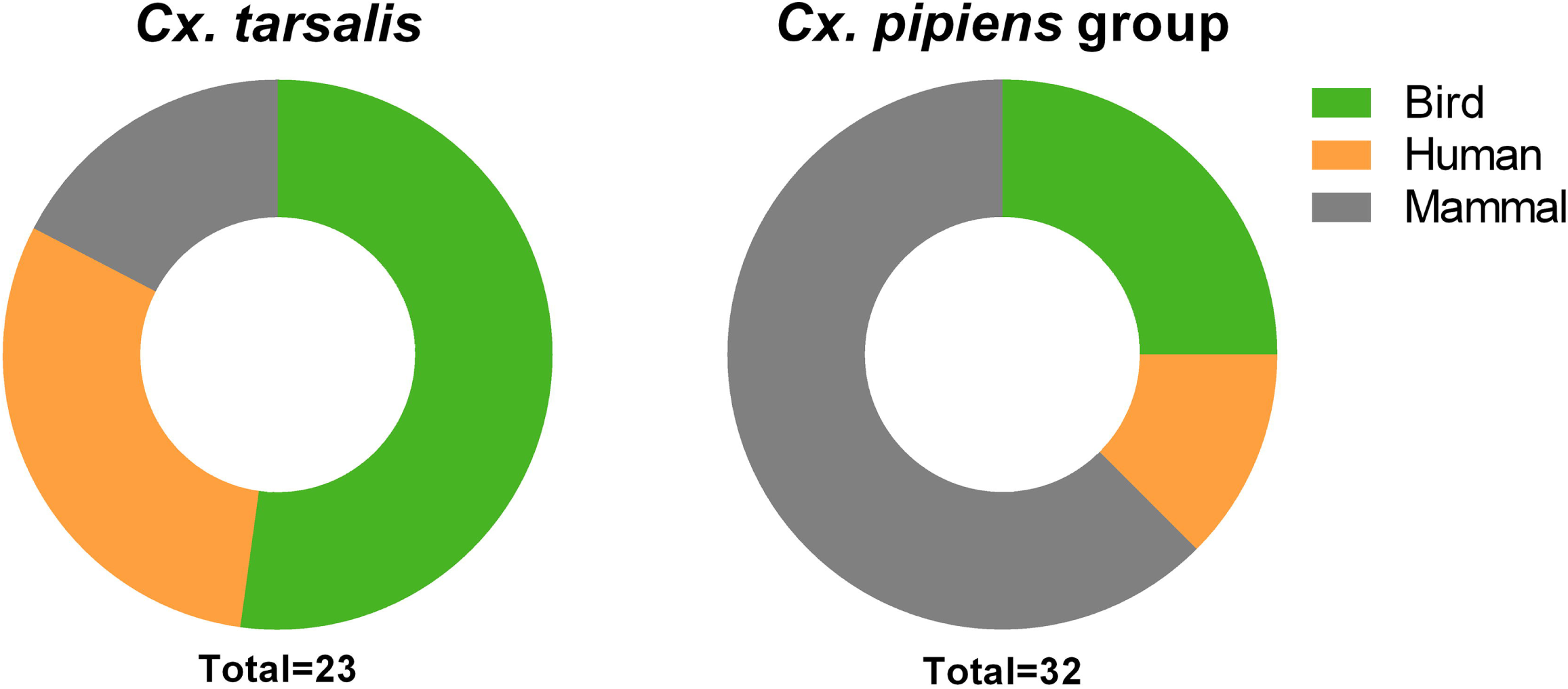
Identification of mosquito blood meals. PCR-based methods were used to identify the blood meal source of collected *Cx. tarsalis* and *Cx. pipiens* group samples. The percentage of blood meals taken respectively from birds, humans, and non-human mammals are displayed for each mosquito species. The total number (n) of identified blood meals is displayed below.

### *Cx. tarsalis* abundance is strongly correlated with human WNV infection

Expanding on initial mosquito abundance data (15), we further examined *Culex* mosquito populations across the state to determine the influence of vector abundance on WNV transmission. Data collected from either CDC or New Jersey Light traps (NJLTs) were used to calculate historical (2002 to 2016) *Culex* mosquito abundance ratios used for comparisons to human WNV incidence on a county-specific level (Fig. 5). These data demonstrate that *Cx. tarsalis* comprises a larger proportion of the total mosquito population in western Iowa than in other regions of the state (Figs. 5A and 5B; 15, 39, 55), and that its relative abundance significantly correlates with human WNV incidence using either CDC (Fig. 5A) or NJLT trap datasets (Fig. 5B). In contrast, *Cx. pipiens* group was widespread throughout the state, without clear associations to human incidence (Figs. 5C and 5D). Together, these data demonstrate that human incidence of WNV was highest in regions where *Cx. tarsalis* was most abundant. When combined with its higher infection rates (Fig. 3) and greater selection for human hosts (Fig. 4), these properties provide strong support towards incriminating *Cx. tarsalis* as the principle vector of the majority of human WNV infections in Iowa.

**Figure 5.**
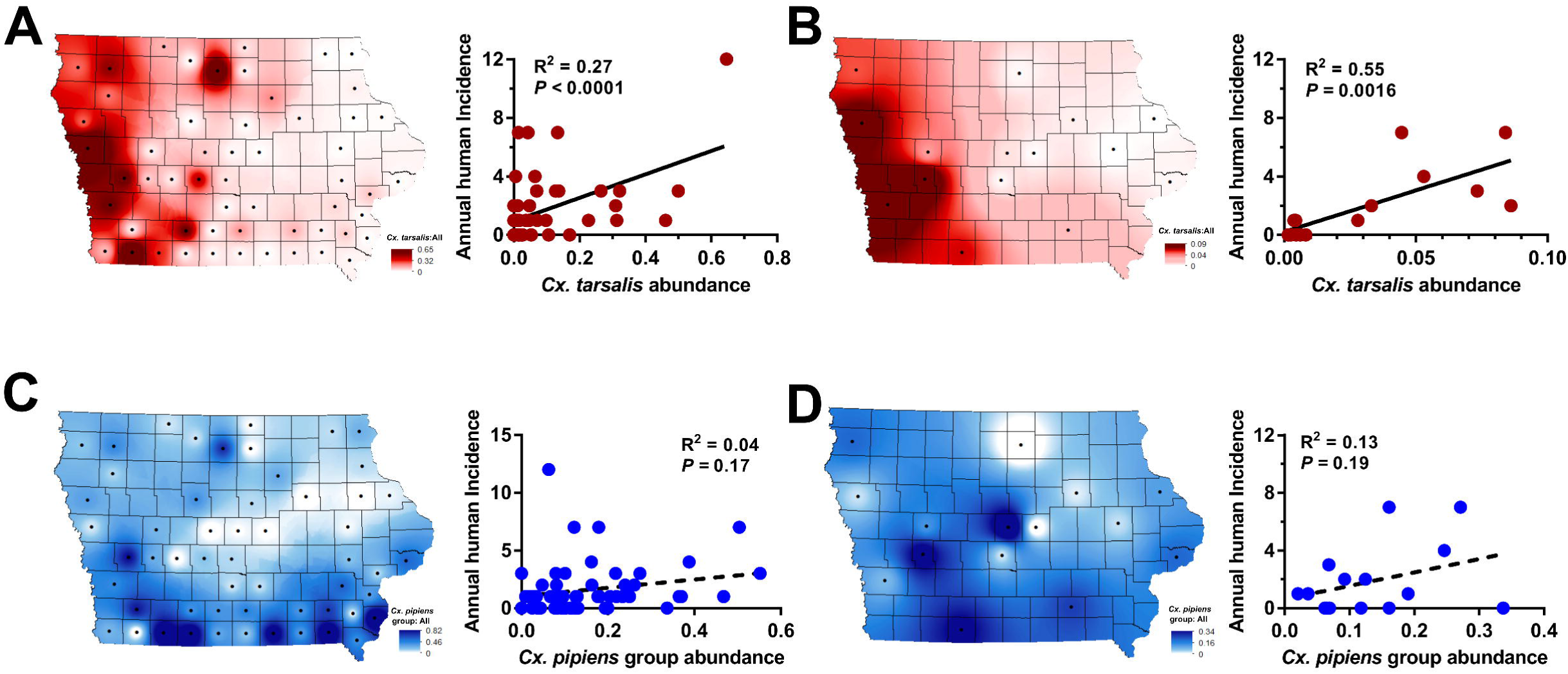
*Cx. tarsalis* abundance significantly correlates with human WNV incidence. Mosquito abundance data collected from CDC traps (A and C) or New Jersey light traps (NJLTs) (B and D) were examined spatially across Iowa. County-specific mosquito datasets were calculated from historical data (2002-2016). Mosquito abundance was measured as a ratio of either *Cx. tarsalis* (A and B) or *Cx. pipiens* group (CPG) (C and D) mosquitoes to the total yield of mosquitoes (all species) collected. Each figure shows a county-specific visual display of mosquito abundance in the state (left) and the relationship of abundance values to human WNV incidence (right). Linear regressions were performed to test the strength of the relationship between mosquito abundance and human WNV incidence, with associated R^2^ and *P*-values displayed for each correlation.

## Discussion

In this study, we provide a comprehensive view of 15 years of WNV transmission in Iowa (2002 to 2016) following its introduction into the state. With the inclusion of infection data from both vertebrate and invertebrate hosts, we provide a unique perspective into the temporal and spatial trends that define WNV transmission in Iowa. We describe a consistent period of WNV transmission in Iowa from late spring to early fall, similar to patterns described in other locations of the Midwest (32, 35) and across the country (9, 27, 56). All host species (human, bird, mosquito) show progressive virus amplification throughout the summer months, with infection rates peaking in late summer for humans and mosquitoes, followed by a dramatic decline with the onset of fall. During this time, WNV remains prevalent in bird populations, which reach peak levels of infection late into the summer and early fall. These transmission dynamics are derived from 15 year averages, yet also display unique inter-annual differences in WNV intensity. The mechanisms driving these infection patterns likely involve interrelated complexities of physiology and behavior that influence interactions between hosts.

Shifts in vector feeding behavior have been previously implicated in driving WNV transmission from an enzootic cycle to human populations (27), yet other studies suggest that little change in feeding preferences occur with the emergence of human cases (31). Therefore, it is unclear how these dynamics may influence human case incidence in Iowa and the greater Upper Midwest region. Additional questions remain regarding the factors that contribute to the dramatic decline of mosquito infection rates and subsequent cessation of human disease cases in early October, despite the presence of WNV in bird populations. This phenomenon is likely a combination of declining temperatures and mosquitoes entering diapause, which would decrease mosquito abundance and biting activity, thereby reducing interactions between vectors and vertebrate hosts.

Similar to initial studies (15), we identify a distinct geographical gradient in the prevalence of WNV across Iowa from east to west. The epidemiological importance of the West is readily evident, where county-level incidence rates in western Iowa were as much as 60-times higher than those in the East. Additional examination of sentinel chicken seroprevalence and mosquito infection rates reinforce western Iowa as a region of epidemiological importance for WNV. Due to the close proximity of Nebraska and South Dakota, which harbor the highest human WNV incidence rates in the country (1), our data imply that the western regions of Iowa may be influenced by similar landscape factors driving WNV transmission as previously suggested (15).

Supported by more than 1.9 million individual mosquito records, we have accumulated a significant body of evidence underscoring the importance of *Cx. tarsalis* in transmitting WNV to humans and in shaping the hyper-endemic region of western Iowa. This is in agreement with previous studies that have proposed important roles for *Cx. tarsalis* in WNV transmission in the Upper Midwest and Northern Great Plains (14, 15, 34, 35). With similar landscape ecology and strong agricultural emphasis across this region, the widespread use of irrigation practices likely provide ideal habitats that enable *Cx. tarsalis* to persist through periods of drought (15, 18). This is in contrast to *Cx. pipiens* group species, which are commonly associated with peridomestic habitats throughout the United States and have been implicated as the primary vector of WNV in urbanized settings (9, 11, 12, 32). While we cannot exclude the involvement and contributions of *Cx. pipiens* group in WNV amplification and human infections, the role of *Cx. pipiens* group mosquitoes is likely that as a secondary vector for WNV transmission in regions where *Cx. tarsalis* is abundant (summarized in Fig. 6). With Iowa serving as the easternmost boundary of *Cx. tarsalis* distribution in North America (57), our data suggest that Iowa serves as an important transition zone, in which *Culex* vector importance shifts along a gradient from *Cx. pipiens* group mosquitoes in the eastern US and Chicago (9, 11, 12, 32) to regions of high *Cx. tarsalis* abundance in Iowa and the Upper Midwest. As a result, highlighting how differences in vector ecology shape distinct epidemiological regions along an ecological gradient in Iowa provides a rare opportunity to examine the species-specific dynamics of WNV transmission in multiple mosquito vectors.

**Figure 6.**
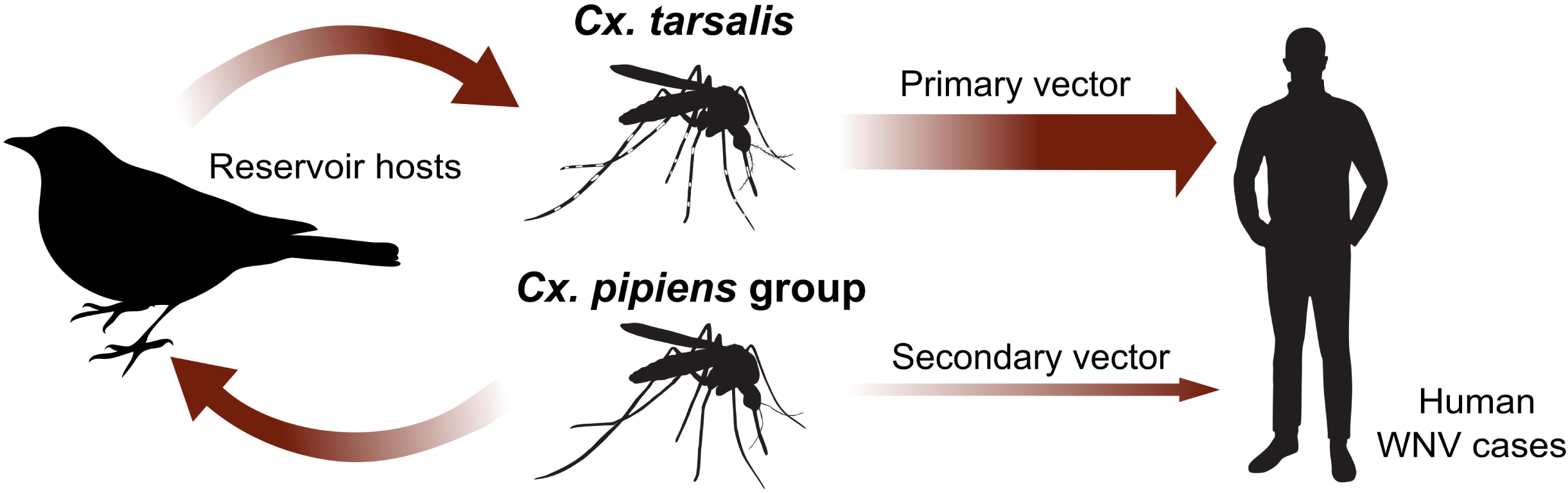
Model of WNV transmission dynamics in Iowa. *Culex* mosquito populations are believed to maintain and amplify WNV in bird reservoirs, with spillover events into human populations primarily occurring through *Cx. tarsalis*. *Cx. pipiens* group mosquitoes are believed to have a secondary role in human cases.

In addition, *Culex pipiens* group mosquitoes display distinct seasonality in their abundance, evidenced by the earlier emergence of *Cx. restuans* before either *Cx. pipiens* or *Cx. salinarius* (20, 58, 59). Previous studies have suggested that *Cx. restuans* may be a comparable or better overall vector of WNV than *Cx. pipiens* (20, 60), such that early season populations of *Cx. restuans* may be integral in establishing WNV amplification in reservoir populations before other vector species promote enzootic spillover into human populations. Collectively referred to as *Cx. pipiens* group in this study, additional efforts should be performed to evaluate the individual contributions of *Cx. pipiens, Cx. restuans*, and *Cx. salinarius* to WNV transmission, especially in the context of their spatiotemporal dynamics.

These differences in vector ecology and abundance are inherently linked to host selection preferences between mosquito species that influence vector competence. *Culex* species have generally been classified as ornithophilic, a preference that promotes frequent interactions with bird reservoirs to acquire and amplify WNV. Several studies have suggested that *Cx. pipiens* group principally feeds on birds (31, 46, 61, 62), while *Cx. tarsalis* is thought to be more opportunistic, feeding on both birds and mammals (53, 62, 63). Our data support previous findings in Iowa (64) suggesting that *Cx. tarsalis* fed on birds more frequently than mosquitoes of the *Cx. pipiens* group.

Moreover, our data suggest that *Cx. pipiens* group fed most commonly on mammals, yet upon further speciation indicate that *Cx. restuans* and *Cx. salinarius* may have stronger preference for non-human mammals while *Cx. pipiens* feed predominantly on birds. *Culex* vectors have been shown to shift increasingly toward mammals through the progression of summer (53, 63–65), a phenomenon that potentially drives epizootic transmission (27), yet the limited scope of our study does not enable temporal analyses. However, it is notable that *Cx. tarsalis* fed on humans more frequently than *Cx. pipiens* group, a potential indicator of greater human host selection by *Cx. tarsalis* that would support a larger role of this vector species in the transmission of WNV to humans. In agreement with these studies, the data presented here support a model in which *Cx. pipiens* group mosquitoes predominantly drive enzootic amplification of WNV in reservoir populations, while *Cx. tarsalis* is more likely to transmit WNV to human populations (Fig. 6).

In summary, our results demonstrate the power and utility of long-term mosquito surveillance to provide a comprehensive understanding of WNV epidemiology and transmission. Integrating human case data, sentinel chicken seroprevalence, and mosquito infection rates, we provide an unprecedented amount of infection data that enable us to define when and where WNV transmission is most active in the state of Iowa. Through these studies, we present strong evidence implicating *Cx. tarsalis* as the primary vector of WNV in Iowa, and identify a unique gradient in vector ecology that shapes virus transmission in Iowa and the Upper Midwest region. With long-term WNV surveillance at the local or state-wide level in many places throughout the country, often with years of unpublished infection and mosquito data, we believe that our study can serve as a template for similar long-term analyses to better understand WNV transmission at the regional or state-wide level. Taken together, our results provide an improved understanding of WNV transmission dynamics in an important epidemiological region of North America.

## Supporting information

## Author Contributions

Conceived and designed the experiments: BMD RCS. Contributed data: BMD KBK EJG ENF WAR LCB RCS. Performed blood meal analysis: EJG ENF. Analyzed the data: BMD KBK EJG RCS. Funding and program oversight: WAR, LCB, RCS. Wrote the initial draft of the manuscript: BMD KBK RCS. Edited the manuscript: BMD LCB RCS. All authors read and approved the final manuscript.

## Acknowledgements

This study would not have been possible without the efforts and data collected by many people that contributed countless hours of labor to the mosquito and WNV surveillance program over the 15 years of this study. We would like to thank collaborators at the State Hygienic Lab, particularly Lucy Desjardin and Thomas Gahan, the local public health partners that assisted in our mosquito surveillance efforts, and the continued support of Ann Garvey and Julie Coughlin from the Iowa Department of Public Health. The authors would also like to thank Katherine Goode for statistical consultation, Hee-Jung Oh for graphic design assistance, and Marilyn O’Hara Ruiz for critical reading of the manuscript. This research was supported by the Iowa State University Agricultural Experiment Station, USDA National Institute of Food and Agriculture, Hatch Project numbers 5033 to WAR (2002-2004), 5111 to LCB (2005-2015) and 101071 to RCS (2016-current), The Midwest Center of Excellence for Vector-Borne Diseases, and Epidemiology and Laboratory Capacity for Infectious Diseases (ELC) Program Components Contract #5887EL11. This publication was supported by Cooperative Agreement #U01 CK000505, funded by the Centers for Disease Control and Prevention. Its contents are solely the responsibility of the authors and do not necessarily represent the official views of the Centers of Disease Control and Prevention or the Department of Health and Human Services.

## References

1. Center for Disease Control and Prevention, West Nile virus. https://www.cdc.gov/westnile/index.html

2. Gray TJ, Webb CE (2014) A review of the epidemiological and clinical aspects of West Nile virus. Int J Gen Med 7:193–203.

3. Ozdenerol E, Taff GN, Akkus C (2013) Exploring the spatio-temporal dynamics of reservoir hosts, vectors, and human hosts of west Nile virus: A review of the recent literature. Int J Environ Res Public Health 10(11):5399–5432.

4. Kramer LD, Styer LM, Ebel GD (2008) A global perspective on the epidemiology of West Nile virus. Annu Rev Entomol 53(1):61–81.

5. Kilpatrick AM (2011) Globalization, Land Use, and the Invasion of West Nile Virus. Science 334(6054):323–327.

6. Kilpatrick AM, Daszak P, Jones MJ, Marra PP, Kramer LD (2006) Host heterogeneity dominates West Nile virus transmission. Proc R Soc B Biol Sci 273(1599):2327–2333.

7. Colpitts TM, Conway MJ, Montgomery RR, Fikrig E (2012) West Nile virus: Biology, transmission, and human infection. Clin Microbiol Rev 25(4):635–648.

8. Andreadis TG (2012) The Contribution of Culex pipiens Complex Mosquitoes to Transmission and Persistence of West Nile Virus in North America. J Am Mosq Control Assoc 28(4s):137–151.

9. Andreadis TG, Anderson JF, Vossbrinck CR, Main AJ (2004) Epidemiology of West Nile Virus in Connecticut: A Five-Year Analysis of Mosquito Data 1999 – 2003. Vector-Borne Zoonotic Dis 4(4):360–378.

10. Ruiz MO, Tedesco C, McTighe TJ, Austin C, Kitron U (2004) Environmental and social determinants of human risk during a West Nile virus outbreak in the greater Chicago area, 2002. Int J Health Geogr 27(2):93–115.

11. Levine RS, Mead DG, Kitron UD (2013) Limited spillover to humans from west nile virus viremic birds in atlanta, georgia. Vector Borne Zoonotic Dis 13(11):812–7.

12. Ruiz MO, Walker ED, Foster ES, Haramis LD, Kitron UD (2007) Association of West Nile virus illness and urban landscapes in Chicago and Detroit. Int J Health Geogr 6(1):10.

13. Resien WK, et al. (2009) Repeated West Nile Virus Epidemic Transmission in Kern County, California, 2004–2007. J Med Entomol 46(1):139–157.

14. Bell JA, Brewer CM, Mickelson NJ, Garman GW, Vaughan JA (2006) West Nile Virus Epizootiology, Central Red River Valley, North Dakota and Minnesota, 2002–2005. Emerg Infect Dis:1245–1247.

15. DeGroote JP, Sugumaran R, Brend SM, Tucker BJ, Bartholomay LC (2008) Landscape, demographic, entomological, and climatic associations with human disease incidence of West Nile virus in the state of Iowa, USA. Int J Health Geogr 7(1):19.

16. Larson SR, DeGroote JP, Bartholomay LC, Sugumaran R (2010) Ecological niche modeling of potential West Nile virus vector mosquito species in Iowa. J insect Sci 10(1):110.

17. Gardner AM, Lampman RL, Muturi EJ (2014) Land Use Patterns and the Risk of West Nile Virus Transmission in Central Illinois. Vector-Borne Zoonotic Dis 14(5):338–345.

18. Schurich J, Kumar S, Eisen L, Moore, Chester G (2014) Modeling Culex tarsalis abundance on the northern Colorado front range using a landscape-level approach. J Am Mosq Control Assoc 30(1):7–20.

19. DeGroote JP, Sugumaran R (2012) National and Regional Associations Between Human West Nile Virus Incidence and Demographic, Landscape, and Land Use Conditions in the Coterminous United States. Vector-Borne Zoonotic Dis 12(8):657–665.

20. Johnson BJ, Robson MG, Fonseca DM (2015) Unexpected spatiotemporal abundance of infected Culex restuans suggest a greater role as a West Nile virus vector for this native species. Infect Genet Evol 31:40–47.

21. Deichmeister JM, Telang A (2011) Abundance of West Nile Virus Mosquito Vectors in Relation to Climate and Landscape Variables Abundance of West Nile virus mosquito vectors in relation to climate. 36(1):75–85.

22. Chuang T-W, Hildreth MB, Vanroekel DL, Wimberly MC (2011) Weather and Land Cover Influences on Mosquito Populations in Sioux Falls, South Dakota. J Med Entomol 48(3):669–679.

23. Turell MJ, Sardelis MR, Dohm DJ, O’Guinn ML (2001) Potential North American vectors of West Nile virus. Ann N Y Acad Sci 951:317–24.

24. Goddard LB, Roth AE, Reisen WK, Scott TW, States U (2002) Vector Competence of California Mosquitoes for. Emerg Infect Dis 8(12):1385–1391.

25. Wimberly MC, Lamsal A, Giacomo P, Chuang TW (2014) Regional variation of climatic influences on West Nile virus outbreaks in the United States. Am J Trop Med Hyg 91(4):677–684.

26. Paull S, et al. (2017) Drought and immunity determine the intensity of West Nile virus epidemics and climate change impacts. Proc R Soc B. doi:10.1098/rspb.2016.2078.

27. Kilpatrick AM, Kramer LD, Jones MJ, Marra PP, Daszak P (2006) West Nile virus epidemics in North America are driven by shifts in mosquito feeding behavior. PLoS Biol 4(4):606–610.

28. Ruiz MO, Tedesco C, McTighe TJ, Austin C, Kitron U (2004) Environmental and social determinants of human risk during a West Nile virus outbreak in the greater Chicago area, 2002. Int J Health Geogr 27(2):93–115.

29. Gu W, Lampman R, Kravasin N, Berry R, Novak R (2006) Spatio-temporal Analyses of West Nile Virus Transmission in Culex Mosquitoes in Northern Illinois, USA, 2004. Vector-Borne Zoonotic Dis 6(1):91–98.

30. Hamer GL, et al. (2008) Culex pipiens (Diptera: Culicidae): A Bridge Vector of West Nile Virus to Humans. J Med Entomol 45(1):125–128.

31. Hamer GL, et al. (2009) Host selection by Culex pipiens mosquitoes and west nile virus amplification. Am J Trop Med Hyg 80(2):268–278.

32. Ruiz MO, et al. (2010) Local impact of temperature and precipitation on West Nile virus infection in Culex species mosquitoes in northeast Illinois, USA. Parasit Vectors 3(1):19.

33. Shand L, et al. (2016) Predicting West Nile Virus Infection Risk from the Synergistic Effects of Rainfall and Temperature. J Med Entomol 53(4):935–944.

34. Bell JA, Mickelson NJ, Vaughan JA (2005) West Nile Virus in Host-Seeking Mosquitoes within a Residential Neighborhood in Grand Forks, North Dakota. Vector Borne Zoonotic Dis 5(4):373–382.

35. Wimberly MC, Giacomo P, Kightlinger L, Hildreth MB (2013) Spatio-temporal epidemiology of human west nile virus disease in South Dakota. Int J Environ Res Public Health 10(11):5584–5602.

36. Chuang TW, Wimberly MC (2012) Remote Sensing of Climatic Anomalies and West Nile Virus Incidence in the Northern Great Plains of the United States. PLoS One 7(10):1–10.

37. Darsie R, Ward R (2005) Identification and geographical distribution of the mosquitoes of North America, North of Mexico. (University Press of Florida).

38. Dunphy BM, Rowley WA, Bartholomay LC (2014) A Taxonomic Checklist of the Mosquitoes of Iowa. J Am Mosq Control Assoc 30(2):119–121.

39. Sucaet Y, Van Hemert J, Tucker B, Bartholomay L (2008) A web-based relational database for monitoring and analyzing mosquito population dynamics. J Med Entomol 45(4):775–84.

40. Biggerstaff B (2006) PooledInfRate, Version 3.0: a Microsoft(r) Excel(r) Add-In to compute prevalence estimates from pooled samples.

41. Malan AK, Stipanovich PJ, Martins TB, Hill HR, Litwin CM (2003) Detection of IgG and IgM to West Nile virus: Development of an immunofluorescence assay. Am J Clin Pathol 119(4):508–515.

42. Post RJ, Flook PK, Millest AL (1993) Methods for the preservation of insects for DNA studies. Biochem Syst Ecol 21(1):85–92.

43. Provost-Javier KN, Chen S, Rasgon JL (2010) Vitellogenin gene expression in autogenous Culex tarsalis. Insect Mol Biol 19(4):423–429.

44. Crabtree M, Savage H, Miller B (1995) Development of a Species-Diagnostic Polymerase Chain Reaction Assay for the Identification of Culex Vectors of St. Louis Encephalitis Virus Bashd on Interspecies Sequence Variation in Ribosomal Dna Spacers. Am J Trop Med Hyg 53(1):105–109.

45. Oshaghi MA, Chavshin AR, Vatandoost H (2006) Analysis of mosquito bloodmeals using RFLP markers. Exp Parasitol 114(4):259–264.

46. Molaei G, Andreadis TG, Armstrong PM, Anderson JF, Vossbrinck CR (2006) Host feeding patterns of Culex mosquitoes and west nile virus transmission, northeastern United States. Emerg Infect Dis 12(3):468–474.

47. Li L, Losser T, Yorke C, Piltner R (2014) Fast inverse distance weighting-based spatiotemporal interpolation: A web-based application of interpolating daily fine particulate matter PM2:5 in the contiguous U.S. using parallel programming and k-d tree. Int J Environ Res Public Health 11(9):9101–9141.

48. Healy J, Reisen W, Kramer V, Barker C (2012) Do surveillance methods provide adequate warning for human infections with West Nile virus. Proc Pap Mosq Vector Control Assoc Calif 80:17–18.

49. Division of Vector-Borne Diseases (2013) West Nile Virus in the United Statesl: Guidelines for Surveillance, Prevention, and Control. http://www.cdc.gov/westnile/resources/pdfs/wnvGuidelines.pdf.

50. Williams GM, Gingrich JB (2007) Comparison of light traps, gravid traps, and resting boxes for West Nile virus surveillance. J Vector Ecol 32(2):285–291.

51. Bolling BG, Moore CG, Anderson SL, Blair CD, Beaty BJ (2007) Entomological Studies Along the Colorado Front Range During a Period of Intense West Nile Virus Activity. J Am Mosq Control Assoc 23(1):37–46.

52. Turell MJ, et al. (2005) An Update on the Potential of North American Mosquitoes (Diptera: Culicidae) to Transmit West Nile Virus. J Med Entomol 42(1):57–62.

53. Kent R, Juliusson L, Weissmann M, Evans S, Komar N (2009) Seasonal Blood-Feeding Behavior of Culex tarsalis (Dipteral: Culicidae) in Weld County, Colorado, 2007. J Med Entomol 46(2):380–390.

54. Kent RJ, Norris DE (2005) Identification of mammalian blood meals in mosquitoes by a multiplexed polymerase chain reaction targeting cytochrome B. Am J Trop Med Hyg 73(2):336–342.

55. Vandyk JK, Rowley WA (1995) Response of Iowa mosquito populations to unusual precipitation patterns as measured by New Jersey light trap collections. J Am Mosq Control Assoc 11(2 Pt 1):200–205.

56. Petersen LR, Brault AC, Nasci RS (2013) West Nile virus: review of the literature. JAMA 310(3):308–15.

57. Venkatesan M, Rasgon JL (2010) Population genetic data suggest a role for mosquito-mediated dispersal of West Nile virus across the western United States. Mol Ecol 19(8):1573–1584.

58. Lee JH, Rowley WA (2000) The abundance and seasonal distribution of Culex mosquitoes in Iowa during 1995-97. J Am Mosq Control Assoc 16(4):275–278.

59. Kunkel KE, Novak RJ, Lampman RL, Gu W (2006) Modeling the impact of variable climatic factors on the crossover of Culex restauns and Culex pipiens (Diptera: Culicidae), vectors of West Nile virus in Illinois. Am J Trop Med Hyg 74(1):168–173.

60. Ebel GD, Rochlin I, Longacker J, Kramer LD (2005) Culex restuans (Diptera: Culicidae) relative abundance and vector competence for West Nile Virus. J Med Entomol 42(5):838–843.

61. Montgomery MJ, Thiemann T, Macedo P, Brown DA, Scott TW (2011) Blood-Feeding Patterns of the Culex pipiens Complex in Sacramento and Yolo Counties, California. J Med Entomol 48(2):398–404.

62. Thiemann TC, et al. (2012) Spatial variation in host feeding patterns of Culex tarsalis and the Culex pipiens complex (Diptera: Culicidae) in California. J Med Entomol 49(4):903–16.

63. Tempelis CH, Reeves WC, Bellamy RE, Lofy MF (1965) A three-year study of the feeding habits of Culex tarsalis in Kern County, California. Am J Trop Med Hyg 14(1):170–177.

64. Ritchie SA, Rowley WA (1981) Blood-Feeding Patterns of Iowa Mosquitoes. Mosq News 41(2):271–275.

65. Tempelis CH, Francy DB, Hayes RO, Lofy MF (1967) Variations in feeding patterns of seven culicine mosquitoes on vertebrate hosts in Weld and Larimer Counties, Colorado. Am J Trop Med Hyg 16(1):111–119.

66. Landesman WJ, Allan BF, Langerhans RB, Knight TM, Chase JM (2007) Inter-Annual Associations Between Precipitation and Human Incidence of West Nile Virus in the United States. Vector Borne Zoonotic Disease 7(3): 337–343.

67. Karki S, Westcott NE, Muturi EJ, Brown WM, Ruiz MO (2017) Assessing human risk of illness with West Nile virus mosquito surveillance data to improve public health preparedness. Zoonoses Public Health (May):1–8.

