## Supplementary material for "Long-term surveillance defines spatial and temporal patterns implicating *Culex tarsalis* as the primary vector of West Nile virus in Iowa, USA"

### Supplemental Information

#### Supplemental Tables

**Table S1.** Summary of combined WNV+ pools for *Culex* species in Iowa, 2002-2016.

#### Supplemental Figures

**Figure S1.** Identification of *Culex pipiens* group blood meals by species.

| Species | # Specimens tested | # Pools tested | # pools WNV+ | MIR |
|---|---|---|---|---|
| *Cx. erraticus* | 4585 | 283 | 6 | 1.31 |
| *Cx. pipiens* group | 129763 | 4888 | 241 | 1.86 |
| *Cx. tarsalis* | 22863 | 731 | 46 | 2.01 |

**Table S1. Summary of combined WNV+ pools for *Culex* species in Iowa, 2002-2016.** Mosquito number, pools tested, and WNV+ pools for each *Culex* species in Iowa. Minimum infection rate (MIR) was calculated as a bias-corrected maximum likelihood estimate.

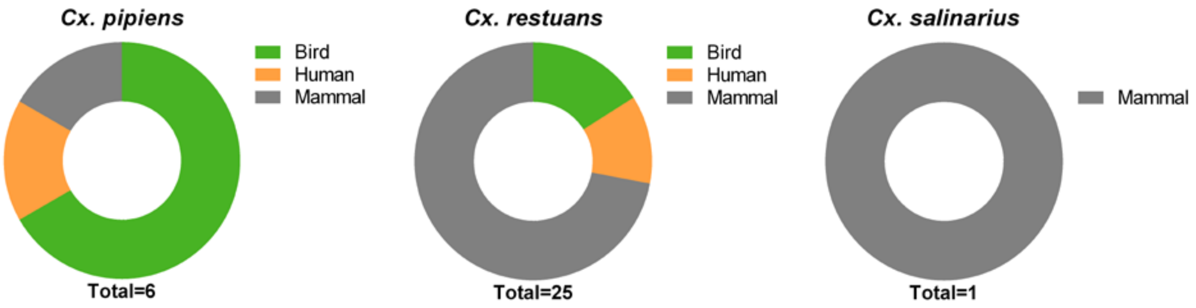


**Figure S1. Identification of *Culex pipiens* group blood meals by species.**

*Culex pipiens* group mosquito samples in which the blood meal source was identified were further speciated to confirm if the sample was *Cx. pipiens*, *Cx. restuans*, or *Cx. salinarius*. The percentage of blood meals taken respectively from birds, humans, and non-human mammals are displayed for each mosquito species. The total number (n) of identified blood meals is displayed below.
